# Surface functionalization of small extracellular vesicles derived from Caco-2 and HEK293T cells in the neutralization of Shiga toxin 1 subunit B

**DOI:** 10.64898/2026.04.23.720326

**Authors:** Krzysztof Mikolajczyk, Anna Bereznicka, Liliana Czernek, Alice Gualerzi, Luana Forleo, Marzia Bedoni, Norbert Lodej, Pawel Migdal

**Author notes:** Corresponding Author: Krzysztof Mikolajczyk, 12 Rudolfa Weigla St., 53-114, Wroclaw, Poland.

## Abstract

Shiga toxins (Stx) are key virulence factors of Shiga toxin-producing *Escherichia coli* (STEC), which are responsible for severe foodborne infections that can progress to hemolytic-uremic syndrome (HUS). Currently, no specific anti-toxin therapies are available. In this study, we devised a glycoengineering strategy utilizing Functional-Spacer-Lipid (FSL) conjugates to create small extracellular vesicles (sEVs)-based decoy receptors for Shiga toxin type 1 (Stx1). sEVs isolated from human Caco-2 and HEK293T cells were functionalized with Gb3 trisaccharide (Galα1→4Galβ1→4Glc)-containing FSL conjugates, yielding Gb3-decorated vesicles, displaying the Galα1→4Gal epitope. Characterization of FSL-modified sEVs confirmed that FSL incorporation did not adversely affect sEV morphology, size distribution, or surface charge. Western blotting and bead-assisted flow cytometry verified the presence of exosomal markers (CD9 and CD63) and the Gb3 epitope on modified vesicles. Gb3-tagged sEVs from both cell types exhibited high specificity in binding Stx1B, while control vesicles carrying Galili epitope (Galα1→3Galβ1→4GlcNAc), lacking Stx1B binding, demonstrated negligible binding. Gb3-expressing Caco-2 cell-based assays revealed that Gb3-decorated sEVs markedly reduced Stx1B binding to Caco-2 cells, indicating effective competition with cellular receptors. Furthermore, glycoengineered sEVs did not impair Caco-2 cell viability at concentrations sufficient for Stx1B sequestration. These findings establish FSL-mediated glycoengineering as a rapid and versatile approach for generating sEV-based decoy receptors that effectively bind Stx1B. Gb3-containing human sEVs may serve as an agent for neutralizing Stx1B and potentially other glycan-binding toxins, supporting the development of promising next-generation anti-toxin therapeutics.

## Introduction

Shiga toxin-producing *Escherichia coli* (STEC) is a group of foodborne pathogens, belonging to multiple serotypes, which are characterized by the release of the Shiga toxins (Stx), its main virulence factor. STEC produces two Stx types: Stx1, identical to the toxin secreted by *Shigella dysenteriae* serotype 1 and more genetically distinct Stx2 ^1^. Stx displays an AB5 structure with one catalytic A domain and a pentamer of B subunits interacting with the Galα1→4Gal epitope, which occurs in Gb3 glycosphingolipid, a major receptor for Stx1 and Stx2 ^2,3^. Recently, a glycoprotein-related Stx receptor, termed P1 glycotope (Galα1→4Galβ1→4GlcNAc-R), capable of binding Stx1 has been identified ^4,5^. Both Stx receptors, Gb3 and P1 glycotope, are synthesized by human α1,4-galactosyltransferase (A4galt) and play essential roles in Stx-mediated disease progression ^3^. Infections by STEC lead to hemorrhagic colitis, which often progresses to hemolytic-uremic syndrome (HUS), a severe complication characterized by thrombocytopenia, anemia and acute kidney failure. Globally, STEC infections are estimated to affect 2.8 million annually, underscoring the epidemiological significance of this pathogen ^6^. In 2023, 30 EU/EEA countries reported 10,901 confirmed cases of STEC infections, compared to 8,565 cases in 2022, resulting in an overall notification rate of 3.2 cases per 100,000 people ^7^.

Currently, no specific therapeutics are recommended for the treatment of Stx-related diseases, and clinical management predominantly relies on supportive care. The use of conventional antimicrobial agents, such as antibiotics, results in the release of toxins following bacterial lysis. Given these limitations, there is an urgent need to explore alternative therapeutic strategies ^8^. Recent efforts have focused on developing Stx neutralizers, such as STARFISH ^9,10^ and Synsorb-Pk ^11–13^, which were primarily designed for local intestinal sequestration of Stx ^8^. In contrast, SUPER TWIG is the only known binder that can eliminate the circulating fraction of Stx ^14^. In these agents, Gb3 trisaccharide analogs were bound to synthetic matrices, such as dendrimers or acrylamide polymers, which may be potentially immunogenic and toxic ^8^. Therefore, next-generation therapeutics should not only exhibit high toxin-neutralizing capacity but also minimize the risk of adverse immune responses, thereby offering improved biocompatibility and therapeutic efficacy.

A particularly promising yet underexplored strategy involves the use of Functional-Spacer-Lipid (FSL) Kode conjugates (Kode Technology, GlycoNZ) to introduce defined functional moieties into lipid bilayers. FSL conjugates possess an amphipathic structure that enables their spontaneous and stable incorporation into biological membranes, including cells, liposomes, and extracellular vesicles (EVs), including small EVs (sEVs). As a result, they decorate the membrane surface with the desired functional group. Structurally, FSLs consist of three components: a functional head group, a spacer and a lipid tail ^15^. The versatility of the functional head group allows for diverse interactions with molecules, including toxins, cells and pathogens, while exhibiting minimal toxicity toward the modified entities ^16^. To date, FSL conjugates have been successfully applied to modify embryonic cells, epithelial/endometrial cells, erythrocytes and pathogens for use in quality control systems and diagnostic panels ^17^ and they have also been explored as potential antiviral agents against HIV ^18,19^.

Here, we proposed the utilization of Gb3 trisaccharide (Galα1→4Galβ1→4Glc) linked to FSL conjugates, incorporated into sEVs to produce Gb3-bearing vesicles. sEVs, naturally occurring EVs with a diameter ranging from approximately 30 to 200 nm, are enclosed by a lipid bilayer membrane, making them an ideal platform for embedding Gb3-FSL conjugates on their surface. sEVs have gained significant attention as vehicles for therapeutic and diagnostic applications due to their inherent biocompatibility, stability in biological fluids, ability to transport bioactive cargo, capacity to traverse biological barriers and facilitate intercellular communication. Notably, sEV-based nanomaterials can be engineered to acquire new functionalities, such as surface-displayed ligands and targeting moieties ^20–22^. The appropriate selection of the cell source can yield the desired therapeutic effect contributed by EVs, such as the anti-inflammatory effect of mesenchymal stromal cell (MSC)-derived EVs, which may be utilized for osteoarthritis treatment ^23^. EVs can also be engineered to enhance their targeting efficiency and therapeutic potential through pre-isolation modification (via genetic engineering, metabolic engineering, or direct manipulation of parent cell membranes) and post-isolation modification (through physical fusion or chemical conjugation), enabling precise regulation of EVs to adapt to specific biological tasks (reviewed in ^22,24^). Modified EVs have demonstrated efficacy in delivering proteins, nucleic acids, and chemotherapeutic agents, with applications covering brain disorders ^25^, tumors ^26^, and liver diseases ^27,28^. Consistent with this growing interest, an increasing number of clinical trials are currently evaluating EV-based therapies across a broad range of indications, including immune-mediated diseases, neurodegenerative disorders, cardiovascular pathologies, and regenerative medicine ^29,30^. To date, more than 300 clinical trials have been registered in the US-NIH clinical trial database (https://clinicaltrials.gov).

In this study, we characterized Gb3 trisaccharide-containing sEVs isolated from human cells as an agent able to bind Stx1B. In our previous work, we demonstrated anti-Stx1B ability of Gb3-contained sEVs isolated from modified Chinese Hamster Ovary Lec2 (CHO-Lec2) cells ^31^. Here, we extend this approach by implementing FSL-based engineering into Caco-2 and HEK293T cells-derived sEVs by incorporating Gb3 trisaccharide molecules into the Caco-2-(exo(Caco)-Gb3-FSL) and HEK293T-derived sEVs (exo(HEK)-Gb3-FSL). The comparative evaluation of two distinct cellular sources for sEV isolation provides insight into how the origin of the parental cell contributes to sEV-mediated anti-Stx activity. Additionally, we investigate the biosafety of glycoengineered sEVs at different Gb3-FSL concentrations. We demonstrate that exo-Gb3-FSL effectively displays the Galα1→4Gal disaccharide and binds Stx1B with high affinity, thereby preventing toxin binding to Caco-2 cells. Collectively, our findings establish a novel glycoengineered method based on FSL incorporation for modifying human sEVs.

## Results

### Preparation and characterization of sEVs from Caco-2 and HEK293T cells

In this study, we isolated sEVs from Caco-2 cells (exo(Caco)) and HEK293T cells (exo(HEK)) using a PEG-based enrichment method with gradient and differential centrifugation steps (Fig 1A). Subsequently, the sEVs underwent glycomodification through incubation with the Gb3-FSL conjugate, resulting in the formation of Caco-2-derived Gb3-FSL-containing vesicles (exo(Caco)-Gb3-FSL) and HEK293T-derived ones (exo(HEK)-Gb3-FSL) (Fig. 1B). As a specificity control, the same sEVs were treated with Galili (Galα1→3Galβ1→4GlcNAc)-FSL, yielding exo(Caco)-Galili-FSL and exo(HEK)-Galili-FSL, respectively. Galili-FSL shares the same FSL lipid backbone as Gb3-FSL, but terminates in a stereochemically distinct epitope (Galα1→4Galβ1 in Gb3-FSL versus Galα1→3Galβ1 in Galili-FSL), not recognized by Stx (Fig. 1B). Particle size and concentration were measured using Zetasizer Ultra Red Label (Malvern Panalytical, UK) and Nanosight NS300 (nanoparticle tracking analysis, NTA, Malvern Panalytical, UK) as previously described ^32,33^. Analysis of dynamic light scattering (DLS) using a Zetasizer Ultra Red revealed distinct size distributions for sEVs derived from Caco-2 and HEK293T cells. Caco-2-derived sEVs, with a dominant peak corresponding to vesicles with a mean hydrodynamic diameter of 101.6 nm (SD = 4.14 nm) and an average particle concentration of 1.187 x 10^9^ particles/ml (SD = 3.3 x 10^8^ particles/ml) (Fig. 1C, upper panel). Conversely, sEVs derived from HEK293T cells displayed three populations, with the predominant fraction characterized by a mean diameter of 255.3 nm (SD = 151.9 nm) and an average particle concentration of 1.039 x 10^8^ particles/ml (SD = 9.267 x 10^7^ particles/ml) (Fig. 1H, upper panel) and upon incorporation of Gb3-FSL conjugates, sEVs derived from Caco-2 cells (exo(Caco)-Gb3-FSL) exhibited three particle populations, with a dominant population showing a mean diameter of 86.41 nm (SD = 6.036 nm) and an average concentration of about 1.294 x 10^9^ particles/ml (SD = 6.922 x 10^8^ particles/ml) (Fig. 1C, bottom panel). In the case of Gb3-FSL-modified sEVs derived from HEK293T cells (exo-(HEK)-Gb3-FSL), two populations were detected, with the major fraction displaying a mean diameter of 116.9 nm (SD = 4.769 nm) and concentration about 2.598 x 10^8^ particles/ml (SD = 2.771 x 10^7^ particles/ml) (Fig. 1H, bottom panel). NTA indicated slightly larger mean diameters, compared to measurements obtained using the Zetasizer Ultra Red, for exo(Caco) (154.0 nm, SD = 5.571 nm), exo(Caco)-Gb3-FSL (156.9 nm, SD = 4.641 nm) and exo(HEK)-Gb3-FSL (143.6 nm, SD = 2.021 nm). In contrast, exo(HEK) exhibited slightly smaller mean particle size (202.8 nm, SD = 7.109 nm) relative to the corresponding Zetasizer Ultra Red measurement. Although the absolute value of particle concentration measured by NTA and Zetasizer Ultra Red Label showed slight differences, no significant variation was observed between modified and non-modified sEVs from both cell sources. Zeta potential measurements revealed that incorporation of Gb3-FSL conjugates resulted in a slight reduction in the negative surface charge of sEVs compared to their unmodified counterparts (Table 1) ^34^. Moreover, the stability and morphology of the sEVs were evaluated using TEM before and after the incorporation of the Gb3-FSL conjugate. TEM analysis of both unmodified and Gb3-FSL-modified sEVs derived from Caco-2 and HEK293T cells revealed intact, spherical vesicles, indicating that Gb3-FSL incorporation did not affect vesicle morphology (Fig. 1D, I) ^33^.

**Fig. 1.**
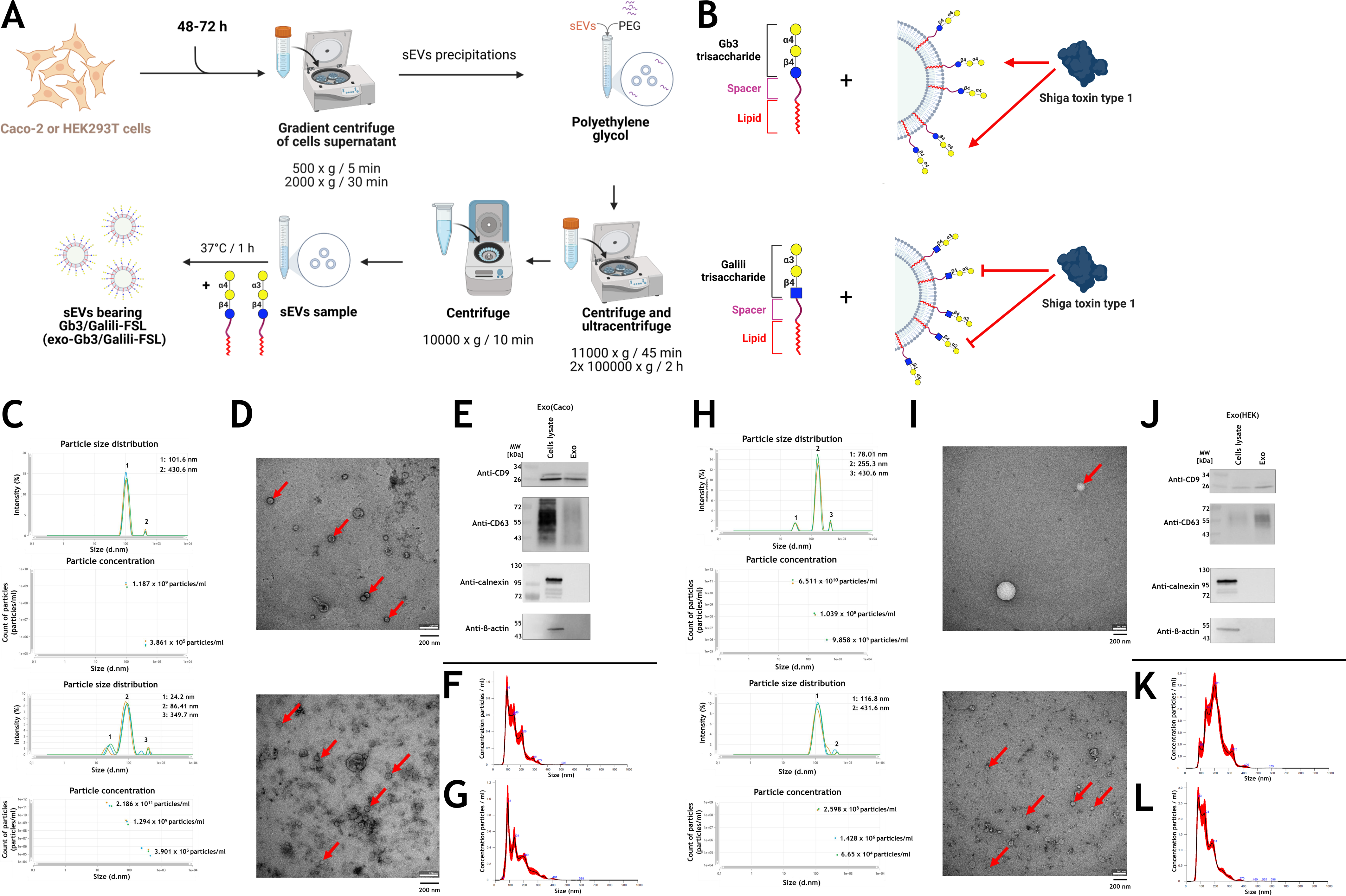
(A) Schematic overview of sEV isolation from Caco-2 and HEK293T cells and their subsequent surface modification using FSL-based conjugates. (B) sEVs isolated from Caco-2 and HEK293T cells were incubated with FSL conjugate bearing either a Gb3 trisaccharide (Galα1→ 4Galβ1→4Glcβ-R) or a Galili trisaccharide (Galα1→3Galβ1→4GlcNAc). Each conjugate consisted of an O(CH_2_)_3_NH spacer and an activated adipate derivative of dioleoylphosphatidylethanolamine lipid. The Gb3 trisaccharide specifically binds the B subunit of Stx1, whereas Stx1B does not recognize the Galili trisaccharide. Characterization of sEVs derived from Caco-2 cells. (C) Representative measurements (the average of at least three repeats) using the Zetasizer Ultra Red showing particle size and concentration of non-modified (upper panel) and modified by Gb3-FSL incorporation (bottom panel), accompanied by TEM visualization of non-modified and modified by Gb3-FSL sEVs from Caco-2 cells (red arrows indicated on representative sEVs) (D). Scale bar for TEM images = 200 nm. (E) Western blotting analysis of non-modified sEVs isolated from Caco-2 cells by incorporation of Gb3-FSL conjugates and corresponding cell lysates, probed with antibodies targeting exosomal markers (CD9 and CD63) and cellular proteins (β-actin and calnexin). (F) NTA analysis of non-modified sEVs from Caco-2 cells. (G) NTA analysis of modified by Gb3-FSL sEVs from Caco-2 cells. The average of the five records was presented. Characterization of sEVs derived from HEK293T cells. (H) Representative measurements (the average of at least three repeats) using the Zetasizer Ultra Red showing particle size and concentration of nonmodified (upper panel) and modified by Gb3-FSL incorporation (bottom panel), accompanied by TEM visualization of non-modified and modified by Gb3-FSL sEVs from HEK293T cells (red arrows indicated on sEVs) (I). Scale bar for TEM images = 200 nm. (J) Western blotting analysis of non-modified sEVs isolated from HEK293T cells by incorporation of Gb3-FSL conjugates and corresponding cell lysates, probed with antibodies targeting exosomal markers (CD9 and CD63) and cellular proteins (β-actin and calnexin). (K) NTA analysis of non-modified sEVs from HEK293T cells. (L) NTA analysis of modified by Gb3-FSL sEVs from HEK293T cells. The average of the five records was presented. Exo(Caco), sEVs derived from Caco-2 cells; Exo(HEK), sEVs derived from HEK293T cells. For statistical analysis, the Kruskal-Wallis nonparametric test, with post-hoc Tukey’s correction, was used. All analyses were performed with GraphPad Prism (GraphPad Software, CA, USA). Statistical significance was assigned to *p*-value <0.05. The part of the figure was created using BioRender.com.

**Table 1.** Size distributions, concentrations and zeta potential of sEVs isolated from Caco-2 and HEK293T cells analyzed by Zetasizer Ultra Red (DLS) and Nanosight NS300 (NTA). The sEVs were treated or not with 100 µM Gb3-FSL conjugate.

| Method | Average size distribution [nm] with SD<sup>1</sup> [nm] | Average particle concentration [particles/ml] with SD<sup>1</sup> [particles/ml] | Polydispersity index with SD<sup>1</sup> | Zeta potential [mV] with SD<sup>1</sup> [mV] |
| --- | --- | --- | --- | --- |
| Exo(Caco) |  |  |  |  |
| DLS | 101.6 (4.14) | 1.187 x 10<sup>9</sup> (3.3 x 10<sup>8</sup>) | 0.199 (0.074) | -19.52 (3.395) |
|  | 430.6 (0) | 3.861 x 10<sup>5</sup> (1.39 x 10<sup>5</sup>) |  |  |
| NTA | 154.0 (2.5) | 8.27 x 10<sup>8</sup> (3.41 x 10<sup>7</sup>) |  |  |
| Exo(Caco)-Gb3-FSL |  |  |  |  |
| DLS | 24.2 (4.167) | 2.186 x 10<sup>11</sup> (1.358 x 10<sup>11</sup>) | 0.263 (0.043) | -28.11 (4.288) |
|  | 86.41 (6.036) | 1.294 x 10<sup>9</sup> (6.922 x 10<sup>8</sup>) |  |  |
|  | 349.7 (89.69) | 3.901 x 10<sup>5</sup> (1.855 x 10<sup>5</sup>) |  |  |
| NTA | 156.9 (2.0) | 7.36 x 10<sup>8</sup> (3.84 x 10<sup>7</sup>) |  |  |
| Exo(HEK) |  |  |  |  |
| DLS | 78.01 (82.74) | 6.511 x 10<sup>10</sup> (6.473 x 10<sup>10</sup>) | 0.223 (0.132) | -35.33 (3.913) |
|  | 255.3 (151.9) | 1.039 x 10<sup>8</sup> (9.267 x 10<sup>7</sup>) |  |  |
|  | 430.6 (0) | 9.858 x 10<sup>5</sup> (5.448 x 10<sup>4</sup>) |  |  |
| NTA | 202.8 (3.2) | 8.45 x 10<sup>8</sup> (3.43 x 10<sup>7</sup>) |  |  |
| Exo(HEK)-Gb3-FSL |  |  |  |  |
| DLS | 116.9 (4.769) | 2.598 x 10<sup>8</sup> (2.771 x 10<sup>7</sup>) | 0.174 (0.030) | -46.77 (6.588) |
|  | 431.6 (42.59) | 7.741 x 10<sup>5</sup> (9.625 x 10<sup>5</sup>) |  |  |
| NTA | 143.1 (0.9) | 2.73 x 10<sup>9</sup> (1.48 x 10<sup>8</sup>) |  |  |

<sup>1</sup> Standard deviation (SD).

Collectively, we successfully isolated sEV subpopulations from both Caco-2 and HEK293T cells, and the incorporation of Gb3-FSL conjugates did not significantly affect sEV size distribution, stability and surface charge.

### Gb3 trisaccharide-containing sEVs from Caco-2 and HEK293T bind Shiga toxin type 1

To date, the use of FSL conjugates embedded in EV lipid bilayers to sequester pathogens or toxins has not been investigated, with the exception of Gb3-containing sEVs derived from CHO-Lec2 cells reported previously for Stx1B binding ^31^. In the current study, we utilized latex bead-assisted flow cytometry to characterize glycoengineered sEVs and to verify the surface display of canonical exosomal marker proteins and the presence of Gb3 epitope.

To verify the exosomal identity and evaluate the purity of sEV preparations derived from Caco-2 and HEK293T cells, we performed western blotting analysis using antibodies that recognized exosomal markers CD9 and CD63. To evaluate potential cellular contamination and the presence of larger vesicular structures, such as microvesicles, the blots were probed with anti-β-actin and anti-calnexin antibodies. Western blotting analysis revealed distinct bands corresponding to CD9 and CD63 in sEV samples (Fig. 1E, J), while no bands were observed for β-actin or calnexin (Fig. 1E, J). In contrast, β-actin and calnexin were readily detected in whole-cell lysates from Caco-2 (Fig. 1E) and HEK293T cells (Fig. 1J), which served as positive controls for cellular proteins ^35,36^.

In all cell-based experiments, we used 100 µM of Gb3-FSL conjugates, which effectively bind anti-P1 (clone 650) and Stx1B, without negatively affecting the integrity of sEVs (irrespective of their origin from Caco-2 or HEK293T cells) (Fig. S1 and Fig. S2). sEVs isolated from Caco-2 and HEK293T cells, modified or not with Gb3-FSL conjugates, were immobilized on 4-µm aldehyde-sulfate latex beads and subsequently stained with antibodies or toxin recognizing exosomal markers (anti-CD9 and anti-CD63) and the Gb3 epitope (anti-P1 (clone 650) and Stx1B-AF488). As a negative control for the latex bead-based flow cytometry assay, the uncoated 4-µm aldehyde-sulfate latex beads were used (Fig. 2A). All bead-bound sEV preparations exhibited surface expression of CD9 and CD63 (Fig. 2B, C). The binding of the anti-P1 (clone 650) antibody, which binds the Gb3 epitope, was detected exclusively on beads coated with Gb3-FSL-modified sEVs derived from either Caco-2 (exo(Caco)-Gb3-FSL) (Fig. 2B) or HEK293T cells (exo(HEK)-Gb3-FSL) (Fig. 2C). The same binding pattern was observed when using Stx1B-AF488 (Fig. 2B, C). We demonstrated approximately 30-fold higher binding capacity of anti-P1 (clone 650) antibody for exo(Caco)-Gb3-FSL vesicles compared to unmodified Caco-2-derived sEVs (exo(Caco)) (Fig. 2B). Stx1B-AF488 binding was 90 times greater for exo(Caco)-Gb3-FSL than for unmodified sEVs (exo(Caco)) (Fig. 2B). For HEK293T-derived vesicles, Gb3-FSL modification resulted in more than a 50-fold increase binding of Stx1B-AF488 (Fig. 2C) and an approximately 240-fold enhacement in anti-P1 (clone 650) antibody binding relative to unmodified control (Fig. 2C). The binding of anti-P1 (clone 650) antibody to unmodified exo(Caco) was approximately 5-fold greater than to unmodified exo(HEK) revealing naturally expression of Gb3 epitope on the vesicle surface (Fig. 2B, C). Exo(Caco)-Galili-FSL and Exo(HEK)-Galili-FSL were used as specificity controls and showed markedly decreased binding capacity of anti-P1 (clone 650) antibody and Stx1B-AF488, confirming the specificity of Gb3-tagged sEVs in Stx1B binding (Fig. 2B, C). Conclusively, Gb3-FSL incorporation resulted in obtaining Gb3-containing sEVs with a high affinity for Stx1B.

**Fig. 2.**
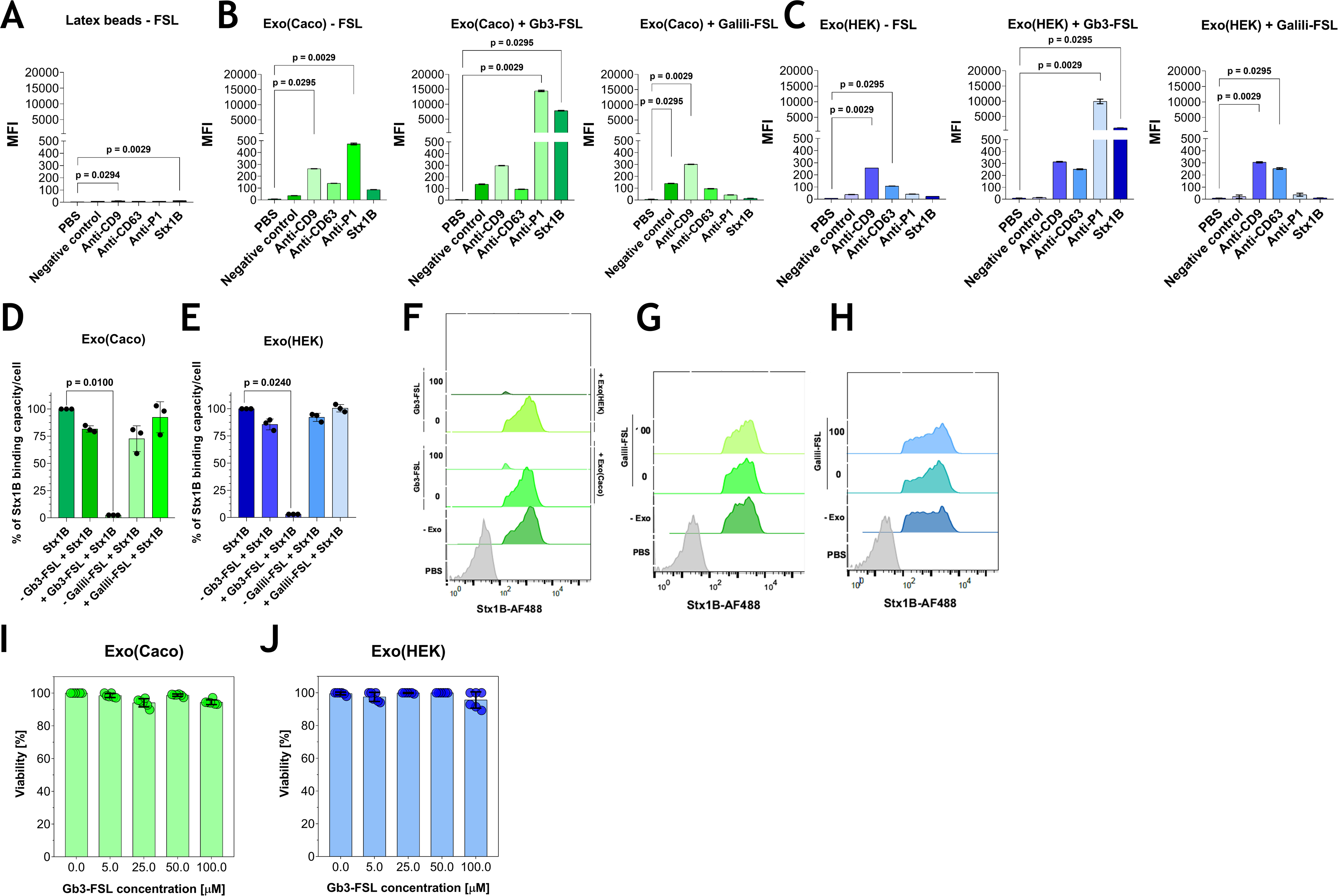
Quantitative flow cytometry analysis of aldehyde/sulfate latex beads with captured sEVs isolated from Caco-2 and HEK293T cells, treated or not with Gb3-or Galili-FSL conjugates. Negative control comprised 4 μm aldehyde/sulfate latex beads without captured sEVs (A), Caco-2 cell-derived sEVs captured on aldehyde/sulfate latex beads (B), and HEK293T cell-derived sEVs captured on aldehyde/sulfate latex beads, stained with anti-CD9-FITC (clone HI9a), anti-CD63-FITC (clone H5C6), anti-P1 (clone 650) antibody and Stx1B-AF488 (C). Average mean fluorescence intensity (MFI) values are shown. Analysis of Stx1B-AF488 binding to Caco-2 cells in the presence or absence of FSL-modified sEVs. The binding of Stx1B-AF488 to Caco-2 cells was evaluated in the presence or absence of sEVs isolated from Caco-2 (D, exo(Caco)) and HEK293T (E, exo(HEK)) cells. For the analysis, data from three repeats were used. Flow cytometry analysis was performed for Stx1B-AF488 coincubated with exo(Caco)-Gb3-FSL and exo(HEK)-Gb3-FSL (F), exo(Caco)-Galili-FSL (G), and exo(HEK)-Galili-FSL (H). Viability assay of Caco-2 cells in the presence of Gb3-FSL-containing sEVs isolated from Caco-2 (I, Exo(Caco)) and HEK293T (J, Exo(HEK)) cells. Gb3-FSL conjugates were incorporated into Caco-2-and HEK293T-derived sEVs at various concentrations (0, 25, 50, 75 and 100 µM). Exo(Caco), sEVs derived from Caco-2 cells; Exo(HEK), sEVs derived from HEK293T cells. For statistical analysis, the Kruskal-Wallis non-parametric test, with post-hoc Tukey’s correction, was used. All analyses were performed with GraphPad Prism (GraphPad Software, CA, USA). Statistical significance was assigned to *p*-value <0.05.

To evaluate the presence of Gb3 epitope and the ability of sEVs to interact with Stx1B, we stained Caco-2-and HEK293T-derived sEVs with and without incorporated Gb3-FSL using the dot-blotting method. This technique preserves vesicle integrity and native membrane organization, unlike SDS-PAGE-based analyses, which may disrupt lipid bilayers and hinder the detection of membrane-associated glycans ^37^, such as the Gb3 trisaccharide. sEV samples were applied to the membrane under non-reducing conditions and, after blocking, probed with anti-P1 (clone 650) antibody and His-tagged Stx1B. All sEV specimens tested positive for CD9 and CD63 (Fig. S3A), confirming their exosomal origin. In contrast, positive staining with anti-P1 (clone 650) antibody and Stx1B was observed exclusively for Gb3-FSL-modified sEVs and non-modified Caco-2-derived ones (Fig. S3A). sEVs modified with the Galili-FSL control did not exhibit any binding (Fig. S3B). These results confirm the successful functionalization of Caco-2-and HEK293T-derived sEVs with Gb3 trisaccharide capable of interaction with Stx1B. Furthermore, spots corresponding to the binding of anti-P1 (clone 650) antibody to non-modified exo(Caco) confirmed native Gb3 epitope expression on these vesicles.

### Gb3 trisaccharide-containing sEVs from Caco-2 and HEK293T can protect the Caco-2 cells from Stx1B binding

We demonstrated that exo(Caco)-Gb3-FSL and exo(HEK)-Gb3-FSL can effectively bind Stx1B. We hypothesized that such glycoengineered vesicles compete with cellular Stx receptors and thereby protect cells from Stx1B binding to the cell surface. To test this hypothesis, we employed Caco-2 cells, a human colorectal adenocarcinoma cell line that naturally expresses Gb3 and binds Stx1B. To confirm the presence of the Gb3 receptor and characterize the Stx1B binding profile, Caco-2 cells were analyzed via flow cytometry following staining with anti-CD77-FITC antibody, which recognizes the Gb3 glycolipid, and fluorescently labeled Stx1B (Stx1B-AF488) (Fig. S4). We demonstrated binding of both antibodies to the cells, confirming the presence of the cell-surface Gb3 epitope (Fig. S4). To further validate receptor specificity, cells were treated with Genz-123346, a selective glucosylceramide synthase inhibitor ^38^, which resulted in a marked reduction in both anti-CD77 and Stx1B-AF488 binding (Fig. S4). These findings confirmed that Caco-2 expresses Gb3 and can bind Stx1B, making it a suitable model for studying Stx-mediated pathogenesis.

Quantitative analysis showed that the incorporation of 100 µM Gb3-FSL into sEVs derived from Caco-2 and HEK293T cells significantly diminished the binding of the Stx1B-AF488 to the cell surface (Fig. 2D-F). Notably, sEVs lacking Gb3-FSL (regardless of their Caco-2 or HEK293T origin) caused a slight reduction in Stx1B-AF488 binding (Fig. 2D-F). This effect may be attributed to the presence of low levels of endogenous Stx receptors, such as Gb3, naturally occurring on cell-derived vesicles, or to nonspecific interactions between the toxin and the vesicle surface. However, the coincubation of Stx1B-AF488 with exo(Caco)-Gb3-FSL and exo(HEK)-Gb3-FSL led to a substantial decrease in the Stx1B binding capacity to Caco-2 cells (Fig. 2D-F), suggesting effective competition of Gb3-expressing sEVs with cellular receptors. To confirm binding specificity, exo(Caco)-Galili-FSL and exo(HEK)-Galili-FSL agents, which lack affinity for Stx1B, were examined. These vesicles did not inhibit Stx1B-AF488 binding to Caco-2 cells (Fig. 2D, E, G, H), confirming the high specificity of exo(Caco)-Gb3-FSL and exo(HEK)-Gb3-FSL for Stx1B and highlighting their potential to protect Stx-sensitive cells by toxin sequestration.

Given the established internalization of sEVs by recipient cells ^39,40^, we proceeded to investigate the uptake of exo(Caco) and exo(HEK) functionalized with Gb3-FSL or not by Caco-2 cells to support the future development of Gb3-FSL-based anti-Stx1 agents. Caco-2-and HEK293T-derived sEVs were labeled with CellTrace™ CFSE dye, a membrane-permeable fluorescent probe that diffuses into vesicles and labels intravesicular proteins. Uptake of CFSE-labeled sEVs by Caco-2 cells was visualized using fluorescence microscopy. Both Caco-2-and HEK293T-derived sEVs were readily detected within the cytoplasm of Caco-2 cells, indicating vesicle internalization (Fig. S5). Caco-2 cells directly labeled with the fluorescent dyes served as a positive control (Fig. S5). The obtained results confirm that sEVs from both Caco-2 and HEK293T cells can be effectively taken up by Caco-2 cells.

### Impact of Gb3 trisaccharide-containing sEVs from Caco-2 and HEK293T on Caco-2 cells viability

To evaluate the biosafety of exo(Caco)-Gb3-FSL and exo(HEK)-Gb3-FSL, cytotoxicity assays were performed using Caco-2 cells. sEVs isolated from Caco-2 or HEK293T cells were functionalized with varying concentrations of Gb3-FSL (0, 5, 25, 50 and 100 µM) in PBS and subsequently added to Caco-2 cells. After 24 h incubation, the cell viability was measured using CellTiter 96 AQueous One Solution Assay. As a positive control, the Caco-2 cells not treated with Gb3-FSL-containing sEVs were used (Fig. 2I, J). Exo(Caco)-Gb3-FSL and exo(HEK)-Gb3-FSL did not affect the viability of Caco-2 cells at any tested concentrations (Fig. 2I, J). Cell viability remained close to 100%, indicating that incorporation of Gb3-FSL into sEVs does not induce cytotoxic effects at concentrations sufficient for toxin sequestration.

## Discussion

The treatment of Stx-mediated diseases is a significant clinical challenge due to the limited efficacy of current antibiotic therapies. Therefore, there is an urgent need for the development of alternative strategies capable of neutralizing the toxin prior to its interaction with host cells. In this study, we demonstrated for the first time that FSL-based glycoengineering can be applied to human cell-derived sEVs to generate multivalent decoy receptors for Stx1B neutralization. By incorporating Gb3 trisaccharide-containing FSL conjugates into sEVs isolated from Caco-2 and HEK293T cells, we created vesicles that display Galα1→4Gal epitope on their surface, which bind Stx1B with high specificity and can inhibit toxin binding to Gb3-expressing Caco-2 cells. Our findings are consistent with previous reports showing that sEVs derived from CHO-Lec2 cells can effectively bind Stx1B ^31^ and that milk-derived sEVs decorated with Gb3 glycolipid derivatives efficiently bind Stx2B ^41^.

sEVs, as carriers of therapeutic molecules, offer various methods for enriching the vesicle membrane with therapeutic molecules. sEVs are characterized by stability in biological fluids, reduced toxicity and immunogenicity relative to synthetic nanoparticles, such as liposomes ^42,43^ or lipid nanoparticles (LNP) ^44^. A promising approach to tackle limits related to EV-based therapeutic pharmacokinetics and tissue tropism is surface functionalization of the exosomal membrane ^45–47^. To date, several methodologies have been used to incorporate functional groups on the vesicle surface, including post-insertion of phospholipid-and cholesterol-related groups. Both methods were initially developed for liposomes ^22,48,49^ and facilitate liposome surface functionalization without the need for organic solvents typically used in conventional liposome preparation. However, the conditions necessary for insertion reaction require either elevated temperatures (for phospholipid-based agent preparation) or reduced temperatures (for cholesterol insertion), which may pose challenges when such prepared liposomes are intended for *in vivo* administration ^49,50^. Recently, macrophage-derived sEVs loaded with paclitaxel and functionalized with aminoethylanisamide-PEG (AA-PEG) were shown to bind specifically to receptors overexpressed on lung cancer cells, highlighting the potential of surface-engineered sEVs for targeted therapeutic delivery ^51^.

Initially, FSL-based conjugates were developed for the attachment of carbohydrate and peptide blood group antigens to the surface of red blood cells (RBCs) using FSL constructs ^52–54^. Here, we present a novel possible application of FSL conjugates, which exhibit high affinity for lipid bilayers, for the functionalization of lipid-enclosed vesicles. Given that a variety of functional groups can be linked to the FSL core and the conjugation process is straightforward and cost-effective, this method offers the rapid production of vesicles with specified surface ligands. A major advantage of the FSL-based method is the possibility of post-isolation vesicle modification, enabling fast and universal surface engineering of sEVs. Unlike genetic engineering strategies, FSL conjugates spontaneously insert into lipid bilayers without interfering with vesicle biogenesis, protein composition, or membrane integrity ^55^. We found that exo(Caco)-Gb3-FSL and exo(HEK)-Gb3-FSL exhibit no adverse alteration in sEV size distribution, morphology or surface charge, indicating that FSL conjugates are non-invasive and safe agents to modify exosomal membrane (Fig. 1C-L).

Analysis of sEVs derived from Caco-2 and HEK293T cells indicates that both cell types are viable sources for Gb3-FSL-related sEV modification and Stx1B sequestration. A low level of binding of the anti-P1 (clone 650) antibody, recognizing the Gb3 epitope, by non-modified sEVs derived from Caco-2 (exo(Caco)), was markedly increased after Gb3-FSL incorporation. However, the non-modified Caco-2 cells-derived sEVs did not show natural affinity to Stx1B (Fig 2B), despite detectable binding of the anti-P1 (clone 650) antibody, indicating the presence of the Gb3 epitope on these vesicles. In our view, the endogenous expression level of Gb3 may be insufficient to facilitate efficient interaction with Stx1B, necessitating further comprehensive studies to assess the membrane localization and organization of native Gb3 within the exosomal membrane. It is well established that variations in the fatty acid composition of Gb3, including modifications such as hydroxylation ^56^, chain length, and degree of unsaturation, can influence the lateral mobility of the lipid in the plasma membrane and alter the conformation of the Gb3-based trisaccharide head group presented on the cell surface ^57–59^. These parameters, along with the surrounding membrane environment, including cholesterol content, play a crucial role in the recognition of host cells by Stx ^2^.

Functionally, Gb3-FSL-containing sEVs effectively competed with endogenous Gb3 receptors localized on Caco-2 cells, resulting in a substantial reduction in Stx1B binding. This mechanism closely mimics the natural interaction between Stx and host receptors, but without triggering cytotoxicity effects. Unlike antibiotic therapy, which may promote toxin release, the proposed decoy receptor-based strategy neutralizes the toxin through direct competition with cellular receptors, resulting in toxin sequestration and functional inactivation. To date, multiple Gb3-based toxin binders have been developed for the treatment of Stx-related diseases (Table 2) ^8^. In comparison to previously developed Stx neutralizers, such as SUPER TWIG, which can eliminate circulating Stx, their synthetic nature and systemic administration requirements may be challenging for clinical translation ^8^. Gb3-FSL-engineered sEVs combine the multivalency necessary for high-affinity binding with the intrinsic biocompatibility of cell-derived EVs. The relatively short half-life of sEVs in human blood minimizes the time of exo-Gb3-FSL conjugates present in the circulation after intravenous administration, resulting in reduced likelihood of potential toxicity and immune response ^43–45^.

**Table 2.** The comparison of developed anti-Stx strategies.

| Anti-Stx strategy | Molecular target/mechanism | Delivery modality | Key advantages | Key limitations | References |
| --- | --- | --- | --- | --- | --- |
| Gb3-synthesis inhibitors (e.g., eliglustat) | - Reduction of cellular Gb3 expression through inhibition of glucosylceramide synthase<br>- Decreased Stx binding and cellular internalization | Small molecule (oral administration) | - Clinically approved<br>- Potential for therapeutic repurposing | - Requires early administration following infection<br>- Limited efficacy once toxin internalization has occurred | 75,76 |
| Gb3-mimetic multivalent receptors (e.g., STARFISH, Synsorb-Pk) | - Multivalent Gb3 analogues competitively bind the B subunit of Stx, preventing receptor engagement | Synthetic polymers or dendrimers (parenteral or oral administration) | - High binding avidity due to multivalency<br>- Direct toxin neutralization | - Limited *in vivo* efficacy<br>- Pharmacokinetic and bioavailability constraints | 9,11–13 |
| Retrograde-transport inhibitors (e.g., tamoxifen) | - Disruption of Stx trafficking from early endosomes to the Golgi apparatus<br>- Redirection of Stx to lysosomal degradation pathways<br>- Induction of toxin destabilization or denaturation | Small-molecule drugs (oral administration) | - Active after toxin binding to the cell surface<br>- Rapid anti-Stx activity | - Requires early intervention after infection<br>- Potential off-target effects<br>- No specific inhibitors are currently approved | 77–81 |
| Monoclonal antibody-based neutralization (e.g., anti-CD77) | - Antibody-mediated binding of circulating Stx, preventing interaction with cellular Gb3 receptors | Human monoclonal antibodies (intravenous administration) | - Well-established production platforms<br>- High target specificity | - High manufacturing cost<br>- Limited tissue penetration<br>- Susceptible to degradation in the gastrointestinal tract (if not administered intravenously) | 82 |
| Gb3-bearing sEVs | - Surface-displayed Gb3 trisaccharides on sEVs sequester the B subunit of Stx, preventing cellular receptor engagement | Surface-engineered extracellular vesicles (intravenous or oral administration) | - Combines multivalent toxin binding with intrinsic biocompatibility<br>- Potential for improved circulation stability | - Requires scalable production and standardization<br>- Heterogeneity of sEV subpopulations | 31,41 |

Generally, the main limitations of sEVs as delivery systems are related to their time-consuming purification and low yield. Nevertheless, new alternatives for naturally released sEVs, such as sEV mimetics ^60,61^, biomimetics ^62^ or cell-derived nanovesicles ^63,64^ are available. Furthermore, sEVs may be modified for cell-specific targeting (increasing the efficiency of the therapy), particularly toward kidney-related cells (a major target for Stx) ^65^. An important consideration for the therapeutic application of Gb3-FSL-modified sEVs is their cellular uptake ^39,66^. Upon the binding of the Gb3-containing sEVs-Stx complex to recipient cells, internalization via endogenous sEV uptake pathways may enhance toxin sequestration and facilitate its intracellular clearance, thereby limiting Stx-related toxicity. However, since Stx holotoxin retains its enzymatically active A subunit after receptor binding, uptake of Gb3-containing-sEVs-Stx complex may provide an alternative route for intracellular toxin delivery. Therefore, although cellular uptake of Gb3-FSL-modified sEVs may improve toxin clearance, their intracellular fate and potential toxicity should be thoroughly investigated to ensure the safety of glycoengineered sEVs-based therapeutics ^66^.

Taken together, we have developed a novel sEV surface-editing strategy based on FSL conjugates for the sequestration of Stx1B. A key advantage of the FSL-based strategy is its ability to rapidly and efficiently decorate sEV surfaces, enabling the targeted delivery of desired ligands within the organism and facilitating the transfer of molecules with therapeutic potential. Looking ahead, glycoengineered sEVs presenting defined carbohydrate epitopes could be adapted to neutralize other carbohydrate-binding pathogens and toxins, such as cholera toxin ^67,68^, pertussis toxin ^69^, influenza virus ^70^ or norovirus ^71^, by acting as decoy sEVs. Such an approach may provide a versatile, biocompatible strategy for targeted sequestration and clearance of diverse pathogens and toxins from the circulation or mucosal surfaces (for instance, in the gastrointestinal tract). We emphasize that FSL-based technology for sEV modification requires further investigations, focusing on obtaining homogeneous EV subpopulations after isolation, minimizing non-EV contaminants, and validating tests using relevant *in vivo* models.

## Experimental Procedures

### Cells and reagents

Human colorectal adenocarcinoma (Caco-2, HTB-37; ATCC, Rockville, MD) and human embryonal kidney (HEK293T, CRL-3216; ATCC, Rockville, MD) were maintained in Dulbecco’s modified Eagle’s medium (DMEM/F12, Thermo Fisher Scientific) supplemented with 10% fetal bovine serum (FBS: Gibco, Gaithersburg, MD) and Pen-Strep (Thermo Fisher Scientific, Inc., Waltham, MA, USA) at 37 °C in a humidified atmosphere with 5% CO_2_. Stx1B-AF488 was conjugated with AlexaFluor 488 according to the Alexa Fluor™ 488 Protein Labelling Kit (Thermo Fisher Scientific, Inc., Waltham, MA, USA). All used antibodies in the study are listed in Table S1.

### sEV isolation from the cell culture supernatant

The sEVs were isolated from Caco-2 and HEK293T cells using polyethene glycol 6000 (PEG6000, Sigma). PEG is commonly used to concentrate virus particles and pelleted vesicles from the cell-derived supernatant ^72^. PEG6000 (final concentration 12%) was combined with filtered water (filtered through a 0,22 µm filter, Merck) and sodium chloride (0,5 M) to make a two-fold concentrated (24% PEG6000 and 1 M NaCl) stock solution.

Vesicle-containing medium from cell culture was centrifuged at 500 g for 5 minutes, followed by 2,000 g for 30 minutes at 4 °C to remove cellular debris and large apoptotic bodies. Once centrifuged, the media was added to an equal volume of a 2× PEG solution at 4 °C, to achieve a desired final PEG concentration (12%). After the 2× PEG solution was added, samples were mixed thoroughly by inversion and incubated at 4 °C overnight (at least 12 hours). The next day, samples were centrifuged at 11000 x g (Sorvall Lynx 6000, Thermo Fisher Scientific, Inc., Waltham, MA, USA) for 45 min at 4 °C. Conical tubes were then decanted. At the same time, the resulting pellet was suspended in 1 mL of particle-free PBS (pH 7.4) and twice ultracentrifuged at 100000 x g for 2 h at 4 °C (Ultracentrifuge, Thermo Fisher Scientific, Inc., Waltham, MA, USA) to wash the particles of contaminating protein and PEG. Finally, the samples were suspended in 250 µl of particle-free PBS (pH 7.4) and stored at 4°C for short-term or at-80°C for long-term storage.

### Preparation and characterization of sEVs bearing Gb3-FSL (exo-Gb3-FSL) or Galili-FSL (exo-Galili-FSL) conjugates

The sEV-containing Gb3-FSL or Galili-FSL conjugates were prepared by coincubation of sEVs (approximately 1 x 10^8^ particles/mL for Caco-2 cells-derived sEVs and 1 x 10^9^ particles/ml for HEK293T cells-derived sEVs) isolated from Caco-2 and HEK293T cells and Gb3-FSL conjugates (stock 1 mM in PBS, Kode Biotech Ltd., Auckland, New Zealand) at 37 °C for 1 h with agitation (350 rpm). To remove unbound Gb3-FSL conjugates, the samples were ultracentrifuged at 100,000×g for 70 min, resuspended in PBS and analyzed using the Zetasizer Ultra Red Label (Malvern Panalytical, UK) to determine particle size, concentration, polydispersity index (PDI) and Zeta potential. Nanoparticle Tracking Analysis (NTA) was used to complement the size distribution and concentration measurements using a Nanosight NS300 system (Malvern Panalytical, UK), an easy-to-use, reproducible platform for nanoparticle characterization (MISEV2023), ^33^. Samples were diluited five fold in fresh PBS prior to analysis. The diluted samples were introduced into the measurement chamber using a syringe pump to ensure a continuous flow during the acquisition. The Brownian motion of the particles was recorded for 60s per measurement, with five independent captures acquired for each sample.

### Flow cytometry of sEV-coated beads

sEVs (100 μl), containing approximately 1 x 10^8^ particles/mL for Caco-2 cells-derived sEVs and 1 x 10^9^ particles/ml for HEK293T cells-derived sEVs, were attached to 4 μm aldehyde/sulfate latex beads (5 µl, Invitrogen, Carlsbad, CA, USA) by mixing in a 100 μl volume of beads for 15 min at room temperature. The suspension was diluted to 1 mL with PBS and incubated overnight at 4°C. The reaction was then stopped with 100 mM glycine. sEV-bound beads were washed in PBS, then washed in 1% BSA/PBS, blocked with 2% BSA/PBS, and stained with anti-CD9-FITC, anti-CD63-FITC, anti-P1 (clone 650) antibodies and Stx1B-AF488. For all samples, 10,000 events of sEVs-containing aldehyde/sulfate latex beads were recorded and then analyzed by Flowing Software (Perttu Terho, University of Turku) and GraphPad Prism (GraphPad Software, CA, USA).

### Transmission electron microscopy (TEM) of sEVs

For TEM analysis, 5 μL of the Caco-2 and HEK293T-derived sEVs suspension in PBS was loaded onto a 400-mesh copper grid with formvar/carbon film and left to absorb for 60 s. Then, the preparation was dried with filter paper. Next, it was negatively stained with 2% uranyl acetate. The images were taken using a JEM-F200 transmission electron microscope (JEOL, Akishima, Japan).

### Western blotting and dot blotting

sEVs and cells (Caco-2 and HEK293T collected after sEV isolation) were suspended in loading buffer. For sEV samples, the buffer did not contain reducing agents (such as β-mercaptoethanol or ditiothreitol), whereas cell samples were supplemented with such agents. The samples were then lysed by incubation at 95°C for 15 minutes. For SDS-PAGE, 20 µg of Caco-2 or HEK293T cell lysates and 25 µl sEV samples, corresponding between 1 x 10^5^ and 1 x 10^6^ particles/ml (depending on sEV samples), were loaded into 4-20% sodium dodecyl sulfate polyacrylamide gels (Thermo Fisher Scientific, Waltham, MA, USA). For western blotting, proteins were transferred to a nitrocellulose membrane (Sigma-Aldrich, St. Louis, MO). The PageRuler Prestained Protein Ladder (Thermo Fisher Scientific, Waltham, MA, USA) was used as a protein standard. For dot blot, 3µl of samples containing Caco-2 or HEK293T cell lysates and exo(Caco/HEK) with or without FSL conjugates were directly applied onto nitrocellulose membranes (Sigma-Aldrich, St. Louis, MO). Cell lysates, for examining the binding of anti-P1 (clone 650) antibody and His-tagged Stx1B, were derived from CHO-Lec2 cells transfected with human *A4GALT* encoding Gb3/CD77 synthase, synthesising Gb3 glycolipid ^73^. Western blotting and dot blotting membranes were processed in parallel using identical conditions.

The membranes were blocked with 5% bovine serum albumin (BSA) suspended in tris-buffered saline solution (TBS), pH 7.2, for 1 h at room temperature. Membranes were then incubated with antibodies recognizing sEV proteins, including anti-CD9 (BioLegend), anti-CD63 (BioLegend), as well as anti-β-actin (Thermo Fisher Scientific, Waltham, MA, USA) and anti-calnexin (Thermo Fisher Scientific, Waltham, MA, USA). After incubation with primary antibodies, membranes were incubated with appropriate biotinylated secondary antibodies, diluted 1:1000 in 1% BSA in TBS and incubated for 1h at 4°C. For the detection of His-tagged Stx1B, a secondary anti-6xHis-tag antibody (Thermo Fisher Scientific) was used, followed by incubation with poly-HRP-streptavidin (Thermo Fisher Scientific). Following three washes with TBS containing 0.05% Tween-20 and distilled water, membranes were incubated with the antibody conjugated to horseradish peroxidase diluted 1:1000 in 1% BSA/TBS for 1 h at room temperature. Protein bands were visualized using SuperSignal™ West Pico PLUS Chemiluminescent Substrate (Thermo Fisher Scientific, Waltham, MA, USA) according to the manufacturer’s instructions. The concentration of cell lysates was determined using the Nanophotometer N60 (Implen, Munich, Germany).

### Flow cytometry of Caco-2 cells

Caco-2 cells were scraped, washed (with PBS), and incubated for 30 min on ice with anti-P1 (clone 650) (all dilutions were done with 1 % BSA in PBS), anti-CD77-FITC antibodies, washed, and incubated with secondary FITC-conjugated antibodies: anti-mouse IgM (only for the sample stained with anti-P1 (clone 650)). To analyze Stx binding, cells were incubated with 1 μg/ml of His-tagged Stx1B, then washed, and incubated with anti-6x-His Tag antibody, followed by washing and incubation with FITC-conjugated anti-mouse IgG. After incubation with FITC-conjugates, the cells were washed, resuspended in 500 μl of cold PBS, and subjected to flow cytometry analysis using an LSRFortessa flow cytometer (BD Biosciences). In total, 1 × 10^5^ events from the gated population were analyzed using FlowJo™ v10.2 Software (BD Life Sciences). To analyze the neutralization ability of FSL-modified sEVs derived from Caco-2 and HEK293T cells, Stx1B-AF488, Gb3-or Galili-FSL conjugates (both at a concentration of 100 µM) were coincubated simultaneously with Caco-2 cells for 1 h and then analyzed using flow cytometry. The Stx1B-AF488 binding capacity per Caco-2 cells was determined using Quantum (Bio-Rad, Hercules, CA, USA) as described in ^74^ or by measuring the median fluorescence intensity. Briefly, binding capacities of Stx1B-AF488 were calculated by interpolation from the curve according to the manufacturer’s protocol and the fluorophore-to-protein molar ratios of FITC-conjugates (AlexaFluor488-and FITC-bound molecules having the same excitation wavelength, thus may be compared with each other). To facilitate visualization, the binding values were normalized and expressed as percentages of total binding per cell. The binding observed in Caco-2 cells incubated with Stx1B-AF488 alone was stated as 100%. Each experiment was repeated three times.

### sEV uptake assay

For immunofluorescence microscopy, cells were seeded at a density of 3 × 10^5^ cells/well in 6 plates and incubated in serum-depleted medium at 37 °C for 4 h with labelled EV derived from either Caco-2 cells (1 x 10^8^ particles/mL) or HEK293T cells (1 x 10^9^ particles/mL). Both Gb3-FSL-containing (100 μM) and unmodified sEVs were tested, while PBS served as the negative control. sEVs were labelled with 25 µM CFSE for 1 h in a thermoshaker protected from light, and excess dye was removed by ultracentrifugation (100000 × g, 2.5 h, 4 °C). The EV pellet was resuspended in filtered PBS before addition to the cells. As a positive control, cells were directly labelled with CFSE (5 µM) for 30 min at 37 °C. As a negative control, CFSE dye alone was subjected to the same ultracentrifugation procedure as EV samples. Cells were grown on cover slips, which were washed with PBS after EV incubation. Phase contrast and fluorescence pictures were taken at 400 x magnification, on a Nikon Eclipse Ti-U microscope equipped with a FITC filter, recorded on a Nikon Digital Sight DS-U2 camera with the help of the NIS-Elements software (instruments and software from Nikon Instruments, Amsterdam, Netherlands).

### Cell viability test

The cell viability was evaluated using the CellTiter 96 AQueous One Solution Assay (Promega, Madison, WI) as described in ^74^. Briefly, Caco-2 cells were seeded (2 × 10^4^ cells/mL) in 96-well plates (Thermo Fisher Scientific, Inc., Waltham, MA, USA) in a complete DMEM/F12 medium (Thermo Fisher Scientific, Inc., Waltham, MA, USA). After 24 h, the medium was replaced with serum-free DMEM/F12 containing varying concentrations (0, 25, 50, 75, 100 µM) of Gb3-FSL-containing sEVs derived from Caco-2 (about 1 x 10^8^ particles/mL) and HEK293T cells (about 1 x 10^9^ particles/mL), then the cells were incubated for an additional 24 h. After treatment with toxins, the morphology of the cells will be assessed and MTS tetrazolium compound (CellTiter 96 AQueous One Solution Assay, Promega, Madison, WI) was added. The plates were incubated in a humidified, CO_2_ atmosphere for 4 h, and the absorbance at 490 nm was recorded using an ELISA plate reader (Clariostar Luminescence Microplate Reader, BMG LABTECH). The background absorbance registered at zero cells/well was subtracted from the data. The absorbance of wells incubated in medium without Gb3-FSL conjugates was taken as 100 % of cell viability. Each experiment was repeated at least three times.

### Statistical analysis

For statistical analysis, the Kruskal-Wallis non-parametric test, with post-hoc Tukey’s correction, was used. All analyses were performed with GraphPad Prism (GraphPad Software, CA, USA). Statistical significance was assigned to *p*-value <0.05.

## Supporting Information

Supporting informations: supplementary figures include optimization of Gb3-FSL concentration for modification of sEVs (Fig. S1 and Fig. S2), dot blot analysis of sEVs (Fig. S3), flow cytometry analysis of Caco-2 cells (Fig. S4) and uptake assay of sEVs (Fig. S5); supplementary table includes used antibodies in the study.

## Author contributions

KM: conceptualization, writing original draft, investigation, data curation, validation and funding acquisition. AB: writing and review original draft, data curation, validation. LC: writing and review original draft, investigation, data curation and validation. AG & LF & MB: investigation, data curation, writing and review original draft. NL & PM.: investigation and data curation.

## Notes

Safety/Hazard Statement. No unexpected or unusually high safety hazards were encountered while performing this work. The authors declare no competing financial interests.

## Acknowledgements

The Authors thanks Prof. Sabina Górska and PhD Agnieszka Razim from the Department of Immunology of Infectious Diseases, Hirszfeld Institute of Immunology and Experimental Therapy, for their assistance and access to the Zetasizer Ultra Red analysis.

## Abbreviations

A4galt: human α1,4-galactosyltransferase
BSA: bovine serum albumin
CFSE: carboxyfluorescein succinimidyl ester
EV: extracellular vesicle
exo(Caco): Caco-2 cells-derived small extracellular vesicles
exo(HEK): HEK293T cells-derived small extracellular vesicles
FACS: flow cytometry
FSL: Function-Lipid-Spacer
Galili: Galα1→3Galβ1→4GlcNAc carbohydrate epitope
Gb3: Galα1→4Galβ1→4Glc carbohydrate epitope
HUS: hemolytic-uremic syndrome
LNP: lipid nanoparticles
NTA: nanoparticle tracking analysis
PBS: phosphate-buffered saline
PEG6000: polyethene glycol 6000 Da
RBC: red blood cells
sEV: small extracellular vesicle
STEC: Shiga toxin-producing *Escherichia coli*
Stx: Shiga toxins

---

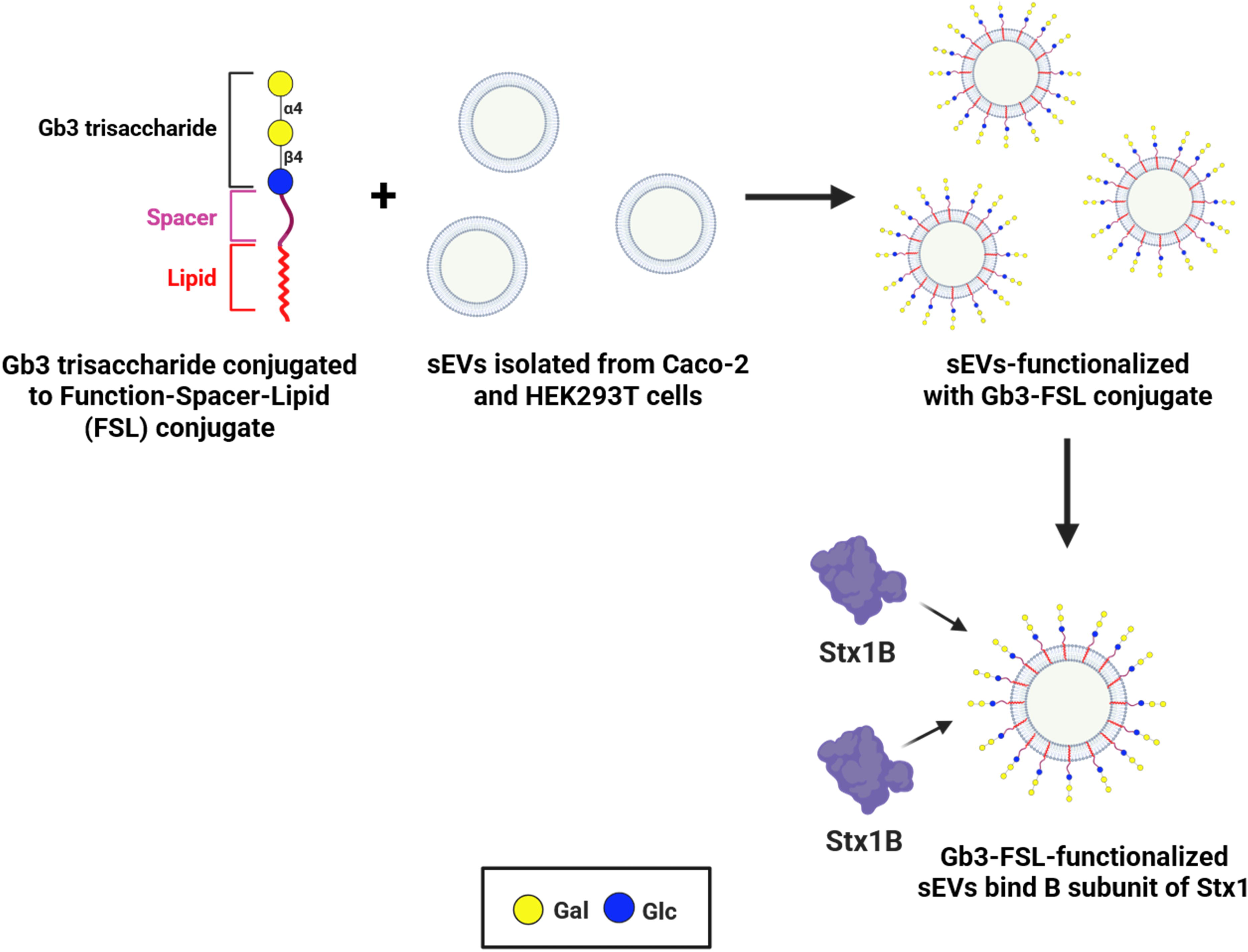

## Notes

### Competing Interest Statement

The authors have declared no competing interest.

### Summary of Updates

Manuscript updated, fugure description updated, TOC figure added.

